# ASAP: A web-based platform for the analysis and interactive visualization of single-cell RNA-seq data

**DOI:** 10.1101/096222

**Authors:** Vincent Gardeux, Fabrice David, Adrian Shajkofci, Petra C Schwalie, Bart Deplancke

## Abstract

**Motivation:** Single-cell RNA-sequencing (scRNA-seq) allows whole transcriptome profiling of thousands of individual cells, enabling the molecular exploration of tissues at the cellular level. Such analytical capacity is of great interest to many research groups in the world, yet, these groups often lack the expertise to handle complex scRNA-seq data sets.

**Results:** We developed a fully integrated, web-based platform aimed at the complete analysis of scRNA-seq data post genome alignment: from the parsing, filtering, and normalization of the input count data files, to the visual representation of the data, identification of cell clusters, differentially expressed genes (including cluster-specific marker genes), and functional gene set enrichment. This Automated Single-cell Analysis Pipeline (ASAP) combines a wide range of commonly used algorithms with sophisticated visualization tools. Compared with existing scRNA-seq analysis platforms, researchers (including those lacking computational expertise) are able to interact with the data in a straightforward fashion and in real time. Furthermore, given the overlap between scRNA-seq and bulk RNA-seq analysis workflows, ASAP should conceptually be broadly applicable to any RNA-seq dataset. As a validation, we demonstrate how we can use ASAP to simply reproduce the results from a single-cell study of 91 mouse cells involving five distinct cell types.

**Availability:** The tool is freely available at http://asap.epfl.ch

## Introduction

Several bioinformatic platforms have been developed that aim to lower the entry point to “-omic” type of analyses (Afgan, et al., 2016; Reich, et al., 2006). The latter include pipelines dedicated to single-cell analyses such as SINCERA (Guo, et al., 2015), SEURAT (Satija, et al., 2015), MAST (Finak, et al., 2015), PAGODA (Fan, et al., 2016) or SC3 (Kiselev, et al., 2016). However, these pipelines are embedded in R which makes them still computationally complex. Furthermore, they only incorporate a restricted set of tools. For example, SC3 has an interactive component but focuses mainly on the clustering part and MAST uses tools that are mainly dedicated to filtering and normalization. Importantly, the available pipelines lack an interactive visualization component as well as integration of a broad range of available single-cell data processing algorithms. In response, several valuable platforms have recently been developed that integrate graphics components. These include SCell (Diaz, et al., 2016), Sincell (Julia, et al., 2015), Fastproject (DeTomaso and Yosef, 2016), or the general RNA-seq analysis platform START (Nelson, et al., 2016). These tools are embedded in standalone applications and cover more comprehensively the whole RNA-seq analysis pipeline, yet, they still lack key features. For example, FastProject performs filtering and visualization but no further analysis. SCell implements RUVg normalization only, and visualization is limited to PCA. Moreover, SCell lacks marker gene identification (based on differential gene expression analysis) or functional gene set enrichment capacities. Finally, all of these pipelines require local installation of the software, which can be time-consuming or even daunting.

To alleviate these constraints, we developed ASAP, a fully integrated, web-based pipeline aimed at the complete analysis of scRNA-seq data post genome alignment. Our choice of rendering ASAP completely web-based was motivated by the fact that less and less users are inclined to install and update manually their tools, which is longer required with web 2.0 software. ASAP allows the user to easily select and compare common, as well as single-cell specific algorithms, and provides an interactive visualization of the results. ASAP support users in the data interpretation process by its fast speed, running the whole analysis pipeline in minutes, and by providing on-the-go visualization, clustering, differential gene expression analysis, and enrichment functionality. ASAP, to our knowledge, is currently the only tool that combines in-depth analysis features and sophisticated visualization for single-cell data in one unique platform.

## Methods

ASAP is a web-based application written in Ruby on Rails. The core structure is completely independent from any currently hosted web application (which are mostly coded in R/Shiny). This effectively makes the platform autonomous and allows the implementation of any tool independent of its source language. Currently, the server runs codes in R, Python and Java, and this process is invisible for the user, who only requires a web browser without prior installation of any development tool. The current list of methods that is included in ASAP is shown in **Figure 1** and detailed in **Supp. Table S1**. Current and past versions are visible in the “about” page of the website (ASAP is versioned according to tool versions).

**Fig. 1.**
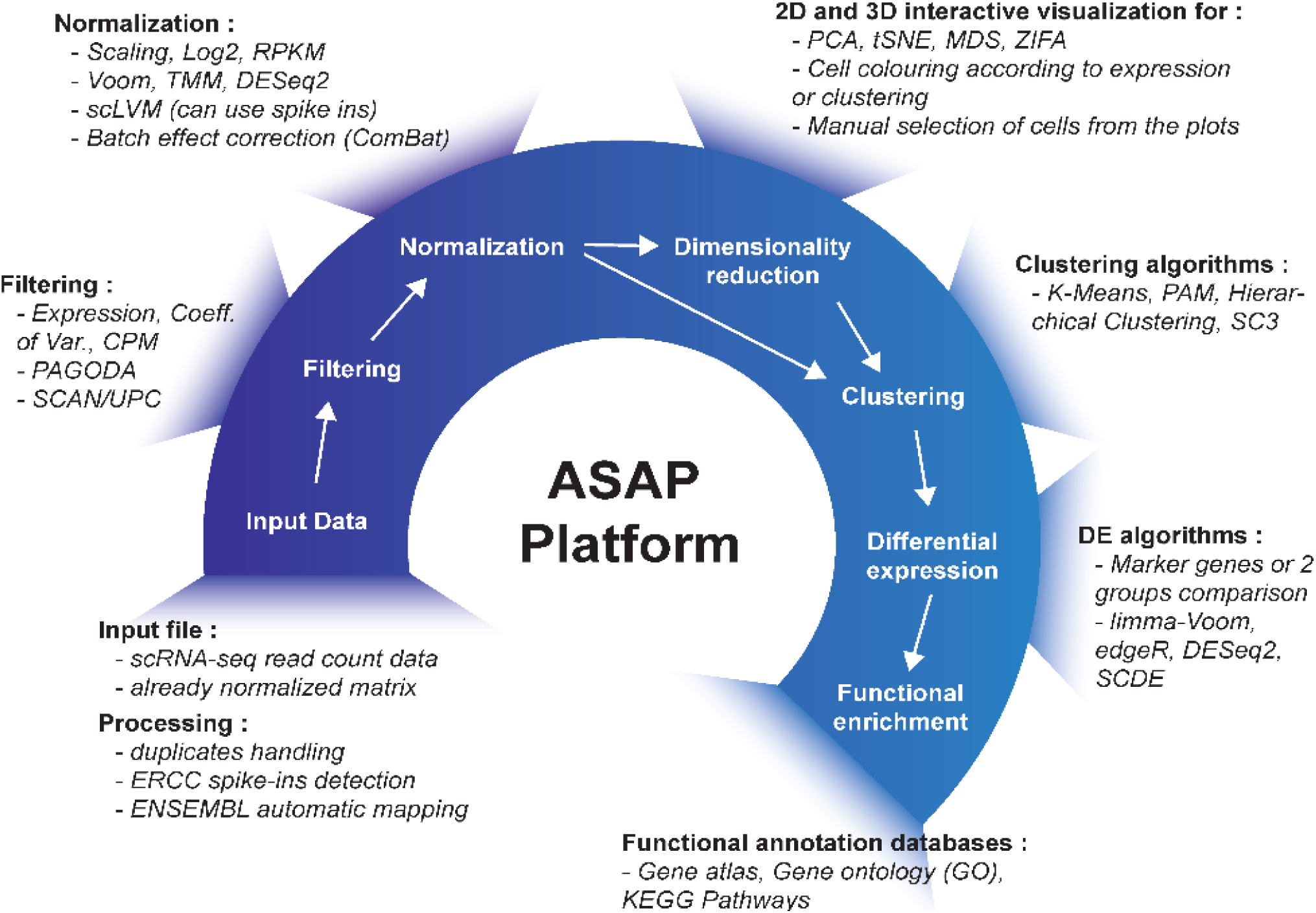
ASAP pipeline. The figure depicts the complete pipeline, including tools, that is implemented in ASAP. The user starts by uploading a count matrix (or normalized matrix) of gene expression after which either the default pipeline or different filtering algorithms can be selected. After the normalization step, the user can apply different dimensionality reductions methods to visualize the data in 2D or 3D. The user can interactively select samples, or run clustering algorithms to perform differential gene expression analysis. Finally, the selected gene list can be analyzed for enrichment in biological modules or pathways such as the Gene Ontology or KEGG. All tools are referenced in **Supp. Table S1**.

The current implementation of ASAP relies on the *delayed::job* framework which automatically creates and queues jobs when the user asks to run a particular method. This allows the application to be perfectly scalable to any IT architecture and prevents major slowdown of the website. Of course, the job execution time scales with the number of users and the host’s computational power. But this will be mainly dependent on the available cores/RAM on the server that hosts ASAP. ASAP has also full compatibility with the last versions of Chrome, Mozilla and Safari. The uploaded user data is protected by an anonymous registration system which keeps the user data private. A sandbox also allows any user to analyze the example project or upload his own data without prior registration. However, the data is destroyed when the user’s session ends.

## Results

As a proof-of-concept, we re-analyzed data from (Dueck, et al., 2015), in which scRNA-seq was used to study gene expression variation across five mouse cell types involving 91 cells. We demonstrate that ASAP is capable of replicating the main findings of this study in minutes in straightforward fashion (**Supp. Figures S1-S14**). We also made these data available as a demo study on the ASAP front page, which is available without registration. It is important to note that, despite the fact that ASAP is primarily dedicated to single-cell analysis, most of the tools can be employed for bulk RNA-seq analysis as well, which makes the pipeline more versatile and universal. ASAP will be further developed as we commit to adding more functionalities such as heatmaps, Venn diagrams, and correlation plots on a continuous basis. We also plan to add an automatic report generation functionality aiming to summarize the employed methods together with figures, version, citation, and parameters. The database for functional enrichment analysis will remain automatically updated through a CRON job, and more databases will be added to cover links to oncogenes, drugs, as well as additional species.

